# So you think you can PLS-DA?

**DOI:** 10.1101/207225

**Authors:** Daniel Ruiz-Perez, Haibin Guan, Purnima Madhivanan, Kalai Mathee, Giri Narasimhan

## Abstract

**Background:** Partial Least-Squares Discriminant Analysis (PLS-DA) is a popular machine learning tool that is gaining increasing attention as a useful feature selector and classifier. In an effort to understand its strengths and weaknesses, we performed a series of experiments with synthetic data and compared its performance to its close relative from which it was initially invented, namely Principal Component Analysis (PCA).

**Results:** We demonstrate that even though PCA ignores the information regarding the class labels of the samples, this unsupervised tool can be remarkably effective as a feature selector. In some cases, it outperforms PLS-DA, which is made aware of the class labels in its input. Our experiments range from looking at the signal-to-noise ratio in the feature selection task, to considering many practical distributions and models encountered when analyzing bioinformatics and clinical data. Other methods were also evaluated. Finally, we analyzed an interesting data set from 396 vaginal microbiome samples where the ground truth for the feature selection was available. All the 3D figures shown in this paper as well as the supplementary ones can be viewed interactively at http://biorg.cs.fiu.edu/plsda

**Conclusions:** Our results highlighted the strengths and weaknesses of PLS-DA in comparison with PCA for different underlying data models.

## Background

### Partial Least-Squares Discriminant Analysis

(PLS-DA) is a multivariate dimensionality-reduction tool [1,2] that has been popular in the field of chemometrics for well over two decades [3], and has been recommended for use in *omics* data analyses. PLS-DA is gaining popularity in metabolomics and in other integrative omics analyses [4–6]. Both chemometrics and omics data sets are characterized by large volume, large number of features, noise and missing data [2,7]. These data sets also often have lot fewer samples than features.

PLS-DA can be thought of as a “supervised” version of *Principal Component Analysis* (PCA) in the sense that it achieves dimensionality reduction but with full awareness of the class labels. Besides its use as for dimensionality-reduction, it can be adapted to be used for feature selection [8] as well as for classification [9–11].

As its popularity grows, it is important to note that its role in discriminant analysis can be easily misused and misinterpreted [2, 12]. Since it is prone to *over-fitting, cross-validation* (CV) is an important step in using PLS-DA as a feature selector, classifier or even just for visualization [13,14].

Furthermore, precious little is known about the performance of PLS-DA for different kinds of data. We use a series of experiments to shed light on the strengths and weaknesses of PLS-DA vis-à-vis PCA, as well as the kinds of distributions where PLS-DA could be useful and where it fares poorly.

The objective of dimensionality-reduction methods such as PCA and PLS-DA is to arrive at a linear transformation that converts the data to a lower dimensional space with as small an error as possible. If we think of the original data matrix to be a collection of *n m*-dimensional vectors (i.e., *X* is a *n × m* matrix), then the above objective can be thought of as that of finding a *m×d* transformation matrix *A* that optimally transforms the data matrix *X* into a collection of *n d*-dimensional vectors *S*. Thus, *S = XA + E*, where *E* is the error matrix. The matrix *S*, whose rows correspond to the transformed vectors, gives *d*-dimensional *scores* for each of the *n* vectors in *X*.

The new features representing the reduced dimensions are referred to as *principal components* (PC). In PCA, the transformation preserves in its first PC as much variance in the original data as possible. On the other hand PLS-DA preserves in its first PC as much covariance as possible between the original data and its labeling. Both can be described as iterative processes where the error term is used to define the next PC. Figure 1 highlights the differences showing an example of a synthetic data set for which the PC chosen by PCA points to the bottom right, while the one chosen by PLS-DA is roughly orthogonal to it pointing to the bottom left.

**Figure 1.**
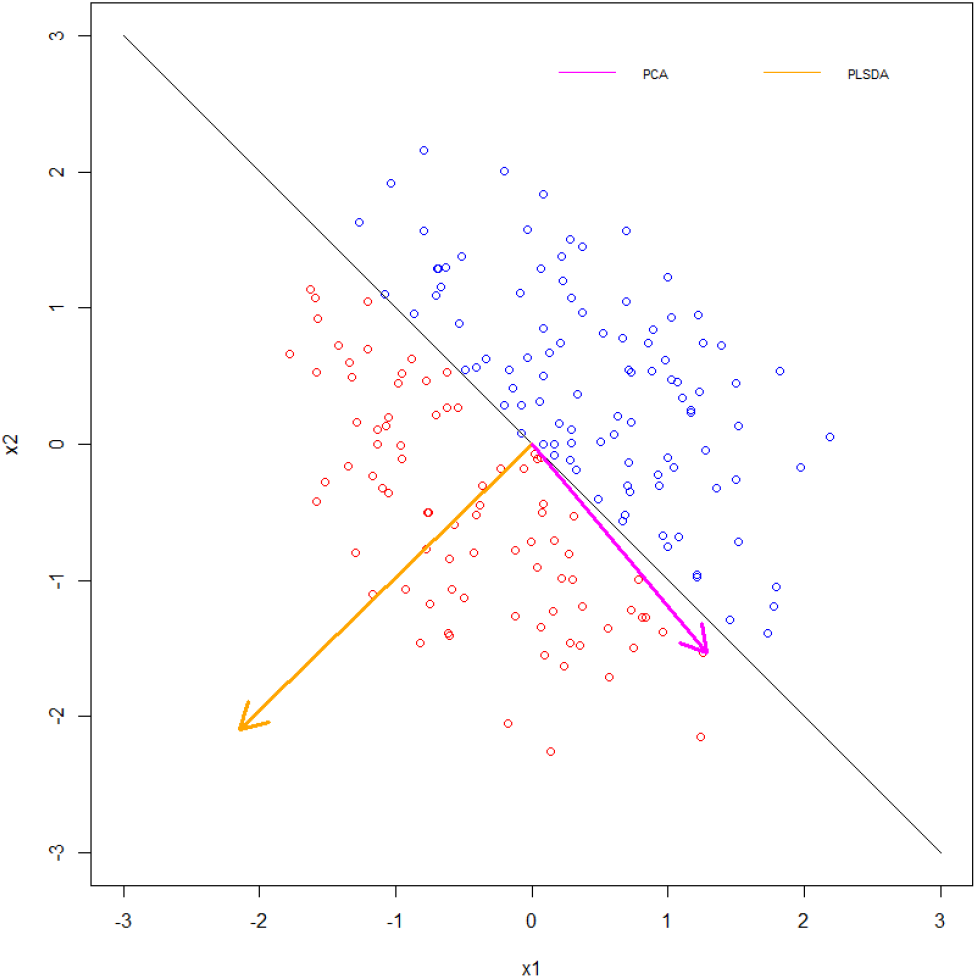
Comparing the first principal component computed by PCA (pink) versus that computed by PLS-DA (orange)

It is also important to note that a higher explained variance or higher correlation for both PCA and PLS-DA doesn’t always mean a better model, even though they are many times linked [14]. The following paragraphs give a more thorough description of the methods and their differences:

### PCA

Informally, the PCA algorithm calculates the first PC along the first eigenvector by minimizing the projection error and then iteratively projects all the points to a subspace orthogonal to the last PC and repeats the process on the projected points. An alternative formulation is that the principal component vectors are given by the eigenvectors of the non-singular portion of the covariance matrix *C* given by:

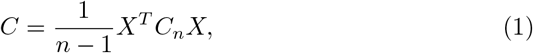

where *C_n_* is the *n × n* centering matrix. The *loading vectors*, denoted by *L*_1_,…, *L_n_*, are given in terms of the eigenvectors, *e*_1_,…,*e_n_* and the eigenvalues, λ_1_,…, λ_*n*_, of *C* as follows:

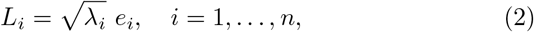

### PLS-DA

In its standard variant the components are required to be orthogonal to each other. In a manner similar to Eq. (1), the first PC of PLS-DA can be formulated as the eigenvectors of the non-singular portion of *C*, given by:

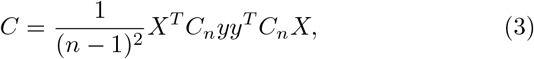

where *y* is the class label vector.

The iterative process computes the *loading vectors a*_1_,…, *a_d_*, which give the importance of each feature in that component. In iteration *h*, it has the following objective:

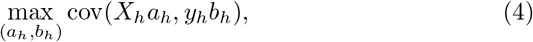

where *b_h_* is the loading for the label vector *y_h_, X*_1_ = *X*, and *X_h_* and *y_h_* are the residual (error) matrices after transforming with the previous *h* − 1 components.

### sPLS-DA

Variant of PLS-DA that makes a *sparsity* assumption, i.e., that only a small number of features are responsible for driving a biological event or effect under study has been devised [15, 16] and shown to be successful with applications where the number of features far outnumber the number of samples [17]. Using lasso penalization, these methods add penalties (*L*_1_ or *L*_0_ norm) to better guide the feature selection and model fitting process and achieve improved classification by allowing to select a subset of the covariates instead of using all of them.

## Methods

In this section, we discuss the aim, design and settings of the experiments.

### Synthetic Data for the Experiments

The following describes a standard experimental setup. Clarifications are provided wherever the experiments differed from this norm. For each of the experiments, labeled synthetic data were generated as follows. The basic input parameters for each experiment are the number of samples *n* and the number of features of each sample *m*. Every data set assumed that there was a *rule* (e.g., a linear inequality), which was a function of some subset of the *m* features (i.e., *signal* features), while the rest were considered as *noise* features. The input parameter also included the rule and consequently the set of signal features. This rule will be considered as the *ground truth*. PLS-DA was then applied to the data set to see how well it performed feature selection or how well it classified. All experiments were executed using PCA and sPLS-DA, where the loading vector is only non-zero for the selected features. Both are available in the *mixOmics* R package [18], which was chosen because it is the implementation most used by biologists and chemists. The noise features of all points are generated from a random distribution that is specified as input to the data generator. The default is assumed to be the uniform distribution. The satisfied rule dictates the generation of the signal features.

### Performance Metrics for the Experiments

As is standard with experiments in machine learning, we evaluated the experiments by computing the following measures: true positives (*tp*), true negatives (*tn*), false positives (*fp*), false negatives (*fn*), precision (*tp* ÷ (*tp + fp*)), and recall (*tp* ÷ (*tp + fn*)). Note that in our case precision and recall are identical. This is because of their formula is the same if *fp = fn*. The data is created with *s* signal features and *s* features are selected. Because *s* is the number of signal features, regardless of whether they were selected or not, *s = tp + fn*. Also, because only *s* features are selected, *s = tp + fp*. Making both equations equal, we get that *fp = fn*.

Since *tn* is large in all our feature extraction experiments, some of the more sophisticated measures are skewed and therefore not useful. For example, the F1 score will be necessarily low, while accuracy and specificity will be extremely high. When the number of noise features is low, precision could be artificially inflated. However, this is not likely in real experiments.

Graphs are shown as 3D plots where the *z* axis represents the performance measure (percentage of signal features in the features marked as important by the tools), and the *x* and *y* axes show relevant parameters of the experiment.

### Experiments varying *n/m*

We first show how the ratio of the number of samples, *n*, to the number of features, *m* affects the apparent performance of PLS-DA and the number of spurious relationships found.

As described earlier, we generated *n* random data points in m-dimensional space (from a uniform distribution) and labeled them randomly. The ratio *n/m* was reduced from 2:1 to 1:2 to 1:20 to 1:200. Given the data set, it is clear that any separation of the data found by any method is merely occurring fortuitously. When we have at least twice as many features as samples, PLS-DA readily finds a hyperplane that perfectly separates both merely by chance. As shown in Figure 2, the two randomly labeled groups of points become increasingly separable. This is explained by the curse of dimensionality, that predicts the sparsity of the data to grow increasingly faster with the number of dimensions.These executions only range in ratios from 2:1 to 1:200. In many current omics data sets, ratios can even exceed 1:1000 (i.e., data sets with 50 samples and 50,000 genes are common). This is one of the reasons of the need of sample size determination when designing an experiment [19].

**Figure 2.**
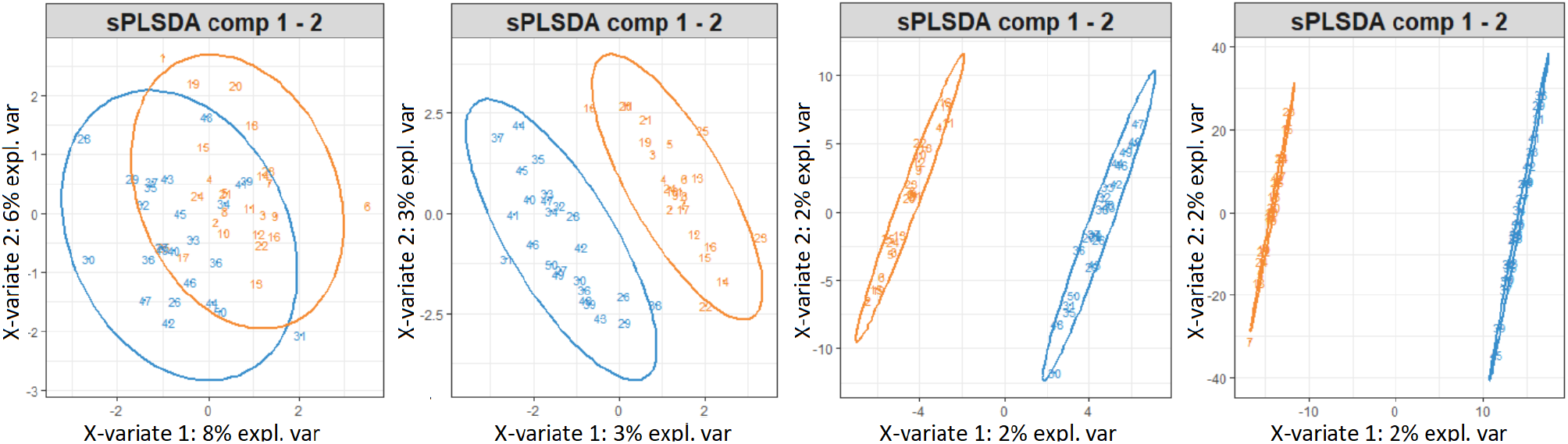
Separability of random points as the ratio of number of samples to features decreases.

If any separating hyperplane is used as a *rule* to discriminate blue points from orange points, then even though the apparent error rate (AE) decreases for this set, its ability to discriminate any new random points will remain dismal [20]. In fact, the CV error rates using 1000 repetitions for the first PC in the four experiments shown in Figure 2 were 0.53, 0.53, 0.5 and 0.48 respectively, showing that even though separability increased, the errors remain unreasonably large. CV errors vary with the seed used to initialize the matrix but the trend remains.

## Results

In this section, we discuss a variety of experiments with synthetic and real data that will help us explain the strenghts and weaknesses of PLS-DA vis-’a-vis PCA and other tools.

### Experiments using PLS-DA as a feature selector

We used 3 sets of methods for generating the synthetic points. In the first set, we consider point sets that are linearly separable. In the second data set we assume that the membership of the points in a class is determined by whether selected signal features lie in prespecified ranges. Finally, we perform experiments with clustered points.

### Experiments with Linearly Separable Points

For these experiments we assume that the data consist of a collection of *n* random points with *s* signal features and *m − s* noise features. They are labeled as belonging to one of two classes using a randomly selected linear separator given as a function of only the signal features. The experiments were meant to test the ability of PLS-DA (used as a feature extractor) to correctly identify the signal features. The performance scores shown in Figure 3 were averaged over 100 repeats. Note that The linear model used implements the following rule 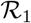, where *C* is a constant set to 0.5:

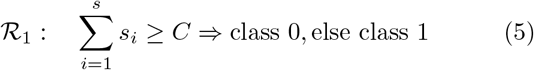

**Figure 3.**
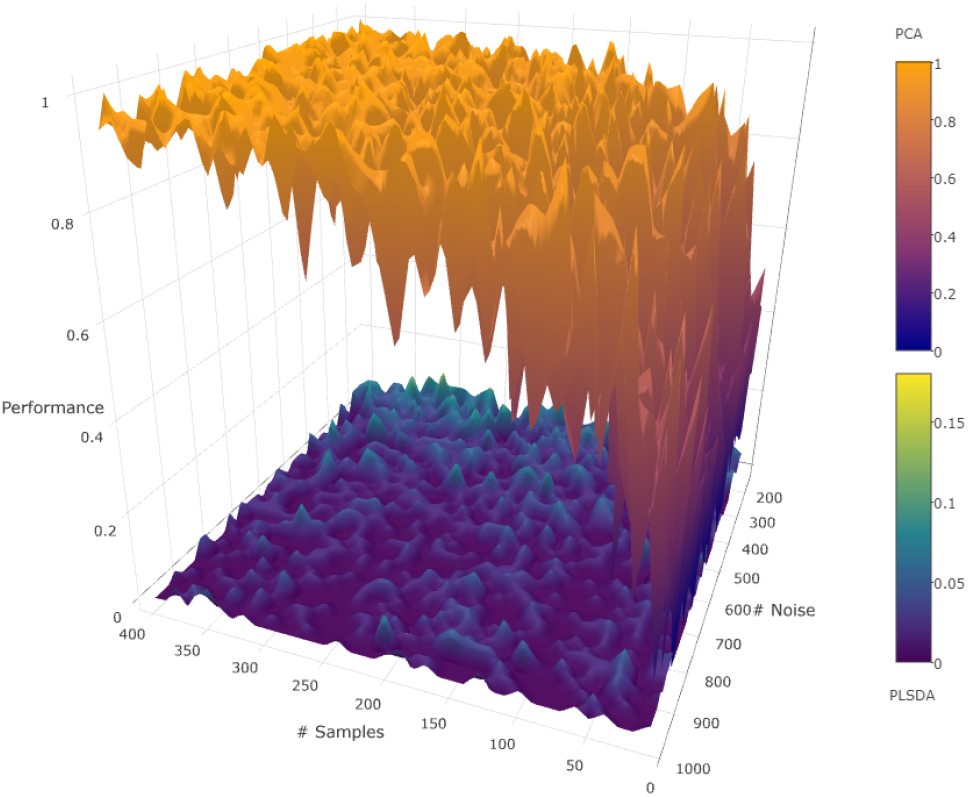
Linear relationship: Samples vs noise.

Two sets of experiments were performed. In the first set, *s* was fixed at 10, but *n* and *m* were varied (see Figure 3). In the second set *n* was fixed at 200, but *s* and *m* were varied (see Additional file 1). PCA consistently outperformed PLS-DA in all these experiments with linear relationships. Also, when the number of samples was increased, the performance of PCA improved, because there is more data from which to learn the relationship. However, it did not help PLS-DA, because the model is not designed to capture this kind of relations. Note that PCA is successful only because the features that are the signal are the only ones correlated.

The *loading vector* is a reflection of what PCA or PLS-DA guessed as the linear relationship between the features. We, therefore, set out to verify how far was the linear relationship that was guessed by the tools used. Even if the tools picked many noise features, we wanted to see how they weighted the noise and signal features they picked. Toward this goal, we ran an extra set of experiments with the model shown above to see if the loading vector from PLS-DA indicated a better performance than what might be suggested by Figure 3. Note that ideally the loading vector should have zeros for the noise features and ones for the signal features. We computed the cosine distance between the loading vector computed in the experiment and the vector reflected by the true relationship. As shown in Additional file 2, we see that the loading vectors of both PCA and PLS-DA failed to reflect the true relationship. These experiments were performed using *n* = 200 averaged over 100 repetitions. Even though PCA successfully selected many of the signal features during feature selection, it was unable to get sufficiently close to the underlying linear relationship, perhaps due to the compositional nature of the signal variables, which gives rise to correlations.

Other experiments carried out with the same results include changing the magnitude of constant in the inequality and changing the relationship from a linear inequality to two linear equalities, i.e., the points lie on two hyperplanes.

### Cluster model

For these experiments, the signal features of the points were generated from a clustered distribution with two clusters separated by a prespecified amount. All noise features were generated from a uniform distribution. The R package *clusterGeneration* was used for this purpose, which also allows control over the separation of the clusters. Cluster separation between the clusters was varied in the range [−0.9,0.9]. Thus when the points are viewed only with the noise features, they appear like a uniform cloud, and when viewed only with the signal features, the members of the two classes are clustered. Note that cluster separation of −0.9 will appear as indistinguishable clusters, while a separation of 0.9 will appear as well-separated clusters. The experiments were executed with *s* = 10, *n* = 200, averaged over 100 repetitions.

The executions with clustered data showed PLS-DA to be clearly superior to PCA. As shown in Figure 4, while it is true that the difference narrows when the number of samples is made very large or the clusters are widely separated (i.e., cleanly separated data),it still remains significant. PLS-DA is able to select the correct hyperplane even with few samples and even when the separation between the clusters is low (values close to 0). PCA needs both an unreasonably large number of samples and very well separated clusters to perform respectably in comparison. However, data with high separation values are embarrassingly simple to analyze with a number of competing methods.

**Figure 4.**
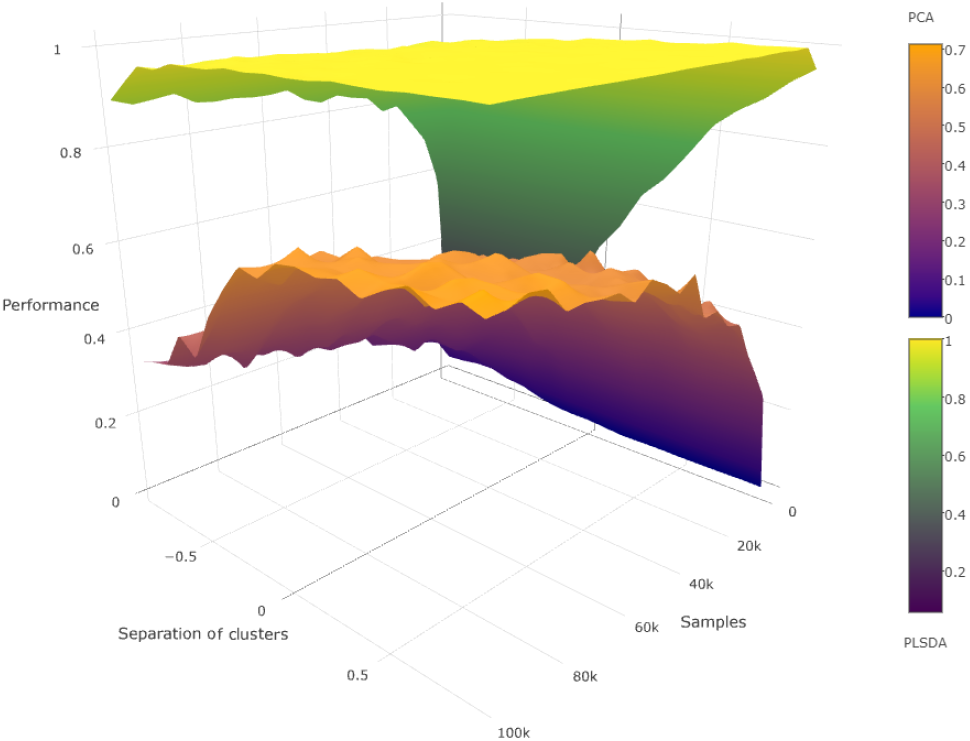
Cluster model: Samples vs cluster separation.

### Interval model

In this set of experiments the rules that determined class membership are often encountered in biological data sets. We used two different methods to generate data from this model. In the first one, we constrained the signal features and in the second we constrained the noise ones. To generate such data sets, members of one class had the constrained features selected uniformly at random from prespecified intervals, while all other features were generated from a uniform distribution in the range [0,1].

We divided the range [0,1] into subintervals of width 1/*p*. Experiments were carried out with *p* = 3, 5 and 10. Depending on the experiment, signal and noise feature were assigned to either a subinterval of width 1/*p* or the entire interval of [0, 1].

The results are shown in Additional file 3. When the signal features are constrained, PLS-DA consistently outperforms PCA. This due to the strong correlation between the signal features for class members that PLS-DA is able to detect. On the other hand, when the noise features are constrained, PCA consistently outperforms PLS-DA. The latter performs poorly when the number of signal features is 1 and *p* = 3, because the distribution of values for the single signal is not very different from the distribution of the noise.

### Experiments as a classifier

Our final set of experiments with synthetic data was to see how PLS-DA fared as a classifier. The following experiments were executed 100 times each, with 10 signal features. For the *cross-validation* error calculation, 5 folds and 10 repetitions were used. In all of the experiments there is a correspondence between a high performance as feature selector and a low CV error.

As shown in Additional file 4a for the linear relationship model, its performance is no better than chance for a 2-class experiment. This corroborates the poor performance of PLS-DA as a feature selector for this model.

For the results with the cluster model shown in Additional file 4b, the CV error is almost 0 in every case, except when the number of samples is low, which is again consistent with what we saw in the feature selection experiments. The performance gets noticeably worse when, in addition to a low number of samples, the number of noise features is large. This is because the signal is hidden among many irrelevant features, something that one has come to expect with all machine learning algorithms. Additional files 4c and 4d show the results for the interval model. As in the case of the feature selection experiments, both versions performed roughly the same, classifying much better than chance and having their best performance when the number of samples was large and the number of noise features was low, as expected.

### Comparisons with other methods

To compare the PLS-DA with other known feature selectors, we applied 3 more methods to the previous data models: *Independent Component Analysis* (ICA), as a feature extraction method that transforms the input signals into the independent sources [21]. *Sparse Principal Component Analysis* (SPCA) via regularized *Singular Value Decomposition* (SVD) [22] was built by adding sparsity constraint. *Regularized Linear Discriminant Analysis* (RLDA) was computed by using *L*_2_ regularization to stabilize eigendecomposition in LDA [23].

We found that PCA based algorithms (PCA and SPCA) have similar overall performance among the three experiments. The same happens with LDA based models (RLDA and sPLS-DA).

As Figure 5, and Additional files 5 and 6 show, PLS-DA, ICA and RLDA are not able to detect linear relationships, while SPCA and PCA are. For the interval model with *p* = 3, either to constrain signal or noise doesn’t seem to change the behavior of the LDA based models, being outperformed by PCA when noise is constrained as shown Additional file 7 and Additional file 8. The performance of every method except for ICA goes down as *s* becomes small.The performance of ICA depends on the number of noise features for both the interval and linear models. In the cluster model experiment as shown in Additional file 9, SPCA performs better than PCA as the separation between the cluster gets higher. The separation between the cluster doesn’t affects the performance of ICA, that stays near 0. RLDA’s and PLS-DA’s performance excel, with similar behavior.

**Figure 5.**
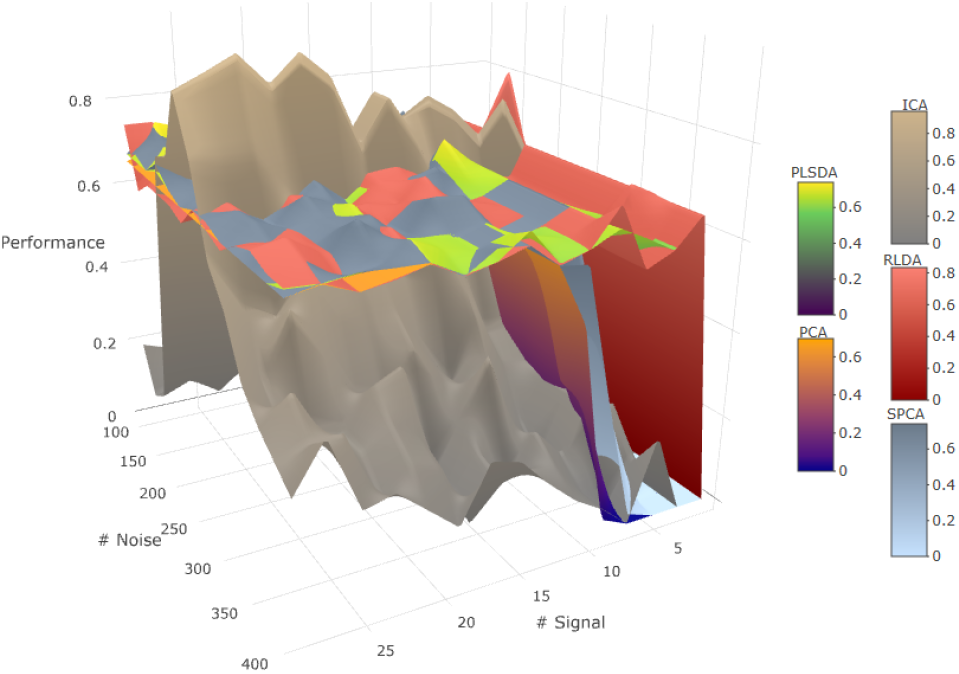
Interval signal constrained for all methods.

### Novel analysis of a real dataset

Bacterial Vaginosis (BV) is the most common form of vaginitis affecting a large number of women across the world [24]. BV is associated with an imbalance of the vaginal flora and damage to the epithelial and mucus layer compromising the body’s intrinsic defense mechanisms. This can result in adverse sequelae and increasing the risk of many STIs [25].

In a landmark paper, human vaginal microbial communities were classified into five community state types (CSTs) [26]. CSTs I, II, III, and V are dominated by different *Lactobacillus* species, whereas CST IV has no specific dominant species and is regarded as the heterogeneous group. While this CST classification has enhanced our understanding of bacterial vaginosis [26–28], a quantitative method to reliably distinguish the CSTs was not available until the development of the specificity aggregation index [29] based on the species specificity [30]. The values of this index range from 0, indicating that the species is absent in that CST to 1, indicating that that OTU is always detected and only detected in that CST.

We used the abundance matrix from [26] (394 samples, 247 OTUs), and with a one vs all approach we devised a simple scheme to differentiate each CST from all of the others using the abundance of each taxon. The importance of each feature given by the specificity index computed in [29] was used as the ground truth. Only the top 10 OTUs for each CST were considered and their importance values were normalized.

Results are summarized in Figure 6. As PLS-DA and PCA return a ranked list of features, a varying threshold on the percentage of features selected is shown on the *X* axis of Figure 6. The *Y* axis represents the sum of the specificity indices achieved by the best features at that cutoff. Note that by just selecting half of the features, a cumulative specificity of 0.9 is achieved by both methods. PLS-DA reaches specificity values over 0.8 with less than 5 features selected, which means that in all of the cases, PLS-DA’s top features are indeed the right set of features. In contrast, PCA’s specificity has a slower growth at the beginning (selects the wrong features), but when half of them are selected both methods achieve the same specificity.

**Figure 6.**
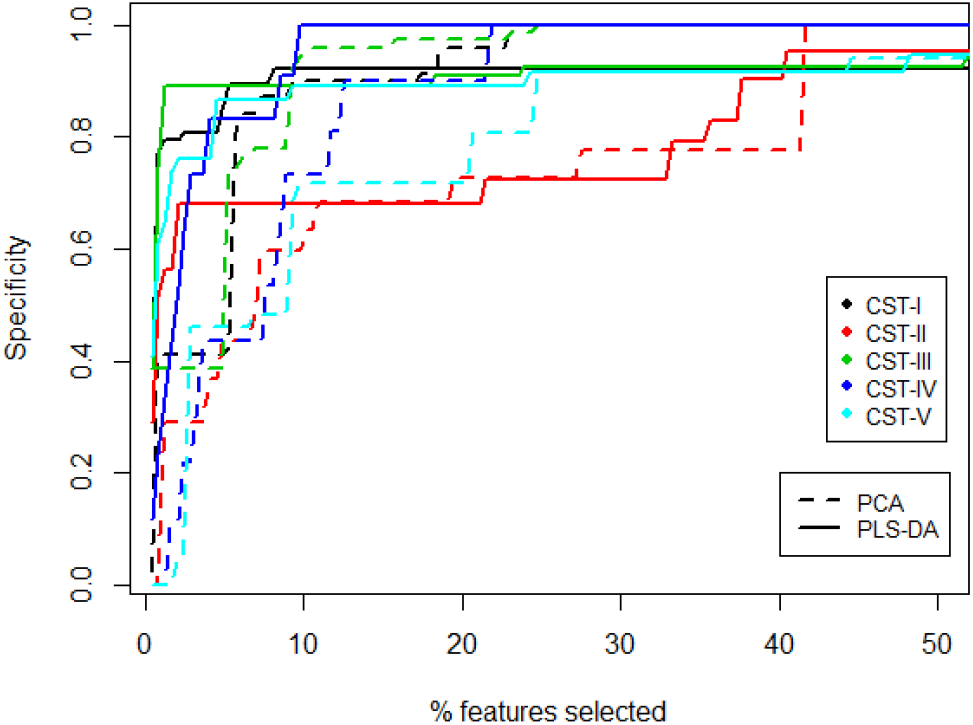
Performance of the features selected by PLS-DA and PCA for different CSTs.

## Discussion

Our work sheds light on the kind of relationships and data models with which PLS-DA can be effective both as a feature selector as well as a classifier. In particular, we claim that when classes are determined by linear or non-linear relationships, PLS-DA provides almost no insight into the data. But it is effective when the classes have a clustered distribution on the signal features, even when these features are hidden among a large number of noise attributes. PLS-DA retains a strong performance also when the classes are contained in n-orthotopes (i.e., rectangular boxes in the subspace of the signal features).

In all of the experiments carried out there was a correspondence between performance of the tools as feature selector and CV error. This reinforces the argument that the CV error is an excellent way to differentiate a good model from a bad one and every paper using PLS-DA must report it to have any validity. Moreover, just-by-chance good behaviors are commonplace when using this tool because the sparsity of the data grows increasingly faster with the number of dimensions and it becomes easier for PLS-DA to find a perfectly separating hyperplane.

Also even though PCA ignores the information regarding the class labels of the samples, it can be remarkably effective as a feature selector for classification problems. In some cases, it outperforms PLS-DA which is made aware of the class labels in its input.

## Conclusions

The obvious conclusion from our experiments is that it is a terrible idea to use PLS-DA blindly with all data sets. In spite of its attractive ability to identify features that can separate the classes, it is clear that any data set with sufficiently large number of features is separable and that most of the separating hyperplanes are just “noise”. Thus using it indiscriminately would turn into a “golden hammer”, i.e., an oft-used, but inappropriate tool. Fortunately, the use of CV would readily point to when it is being used ineffectively.

Our work sheds light on the kind of relationships and data models with which PLS-DA can be effective and should be used both as a feature selector as well as a classifier in the case that the underlying model of the data is known or can be guessed. When it is not possible, one should rely on the CV error and use extreme care when making conclusions and extrapolations.

Also, one should take advantage of the multitude of tools available and use different methods depending on the dataset, as the simple PCA was able to outperform PLS-DA depending on the conditions.

## List of abbreviations

PLS-DA: Partial Least-Squares Discriminant Analysis
PCA: Principal Component Analysis
CV: Cross-Validation
PC: Principal Components
sPLS-DA: Sparse Partial Least-Squares Discriminant Analysis
tp: true positives
tn: true negatives
fp: false positives
fn: false negatives
SPCA: Sparse Principal Component Analysis
ICA: Independent Component Analysis
RLDA: Regularized Linear Discriminant Analysis
SVD: Singular Value Decomposition

## Declarations

### Ethics approval and consent to participate

Not applicable.

### Consent for publication

Not applicable.

### Availability of data and material

All the code that generated and analyzed the datasets can be downloaded from https://github.com/DaniRuizPerez/SoYouThinkYouCanPLS-DA_Public.

All the 3D figures shown in this paper as well as the supplementary ones can be viewed interactively at http://biorg.cs.fiu.edu/plsda. The dataset analyzed are derived from the following published article: [26]

### Author’s contributions

DR-P was the major contributor of this work. DR-P and GN conceived and designed the experiments. DR-P implemented the experiments. DR-P and HG executed the experiments. PM and KM provided the vaginal dataset and helped with the biological analysis. DR-P, GN and HG contributed in writing the manuscript, All authors read and approved the final manuscript.

## Acknowledgments

The authors thank the members of the Bioinformatics Reseach Group (BioRG) for their useful comments during the course of this research.

## Additional Files

Additional file 1: **Figure S1.** Performance for linearly separable points model, varying signal and noise.

Additional file 2: **Figure S2.** Performance for linearly separable points model with the cosine model, varying signal and noise.

Additional file 3: **Figure S3.** Performance table for different configurations of the interval model.

Additional file 4: **Figure S4.** Classification accuracy for the different data models.

Additional file 5: **Figure S5.** Performance for linearly separable points model, varying signal and noise.

Additional file 6: **Figure S6.** Performance for linearly separable points model, varying samples and noise.

Additional file 7: **Figure S7.** Performance of other methods, signal constrained interval with p=3.

Additional file 8: **Figure S8.** Performance of other methods, noise constrained interval with p=3.

Additional file 9: **Figure S9.** Performance of other methods for the cluster model, High number of samples.

